# Steroid hormones sulfatase inactivation extends lifespan and ameliorates age-related diseases

**DOI:** 10.1101/541730

**Authors:** Mercedes M. Pérez-Jiménez, Paula Sansigre, Amador Valladares, Mónica Venegas-Calerón, Alicia Sánchez-García, José M. Monje, Sara Esteban-García, Irene Suárez-Pereira, Javier Vitorica, José Julián Ríos, Marta Artal-Sanz, Ángel M. Carrión, Manuel J. Muñoz

## Abstract

Aging and fertility are two interconnected processes. From invertebrates to mammals, absence of the germline increases longevity by a still not fully understood mechanism. We find that loss of function of *sul-2*, the *Caenorhabditis elegans* steroid sulfatase (STS), raises the pool of sulfated steroid hormones and increases longevity. This increased longevity requires factors involved in germline-mediated longevity (*daf-16*, *daf-12*, *kri-1*, *tcer-1* and *daf-36* genes) and is not additive to the longevity of germline-less mutants. Noteworthy, *sul-2* mutations do not affect fertility. Thus, STS inactivation affects the germline signalling process regulating longevity. Interestingly, *sul-2* is only expressed in sensory neurons, suggesting a regulation of germline longevity by environmental cues. We also demonstrate that treatment with the specific STS inhibitor STX64, reproduces the longevity phenotype of *sul-2* mutants. Remarkably, STS inhibition by either mutation or drug treatment ameliorates protein aggregation diseases in *C. elegans* models of Parkinson, Huntington and Alzheimer, as well as Alzheimer disease in a mammalian model. These results open the possibility of reallocating steroid sulfatase inhibitors for the treatment of aging and aging related diseases.

---

Unravelling new elements that govern the genetic control of aging is key to improve our understanding of this intricate biological process and improve human healthspan. To this aim we isolated *Caenorhabditis elegans* thermotolerant mutants using a previously reported protocol^1^, and identified an allele *pv17* of the *sul-2* gene, which encodes one of the three members of the *C. elegans* sulfatase family^2^. Worms carrying either the isolated point mutation (*pv17*) or the null allele *(gk187)* of *sul-2* live longer than wild type (Fig. 1a-b, Extended Data Fig.1a-c and additional data of all longevity assays in Supplementary Table 1) and enhances the development phenotypes of mutants in the insulin/insulin like growth factor (IGF) receptor *daf-2* (Extended Data Fig. 1d-f), which were used for gene mapping and identification. The *pv17* allele introduces a single amino acid substitution (G46D) resulting in a loss of function phenotype. The curated sequence slightly differs from the one published^3^ (Extended Data Fig. 2).

**Figure 1.**
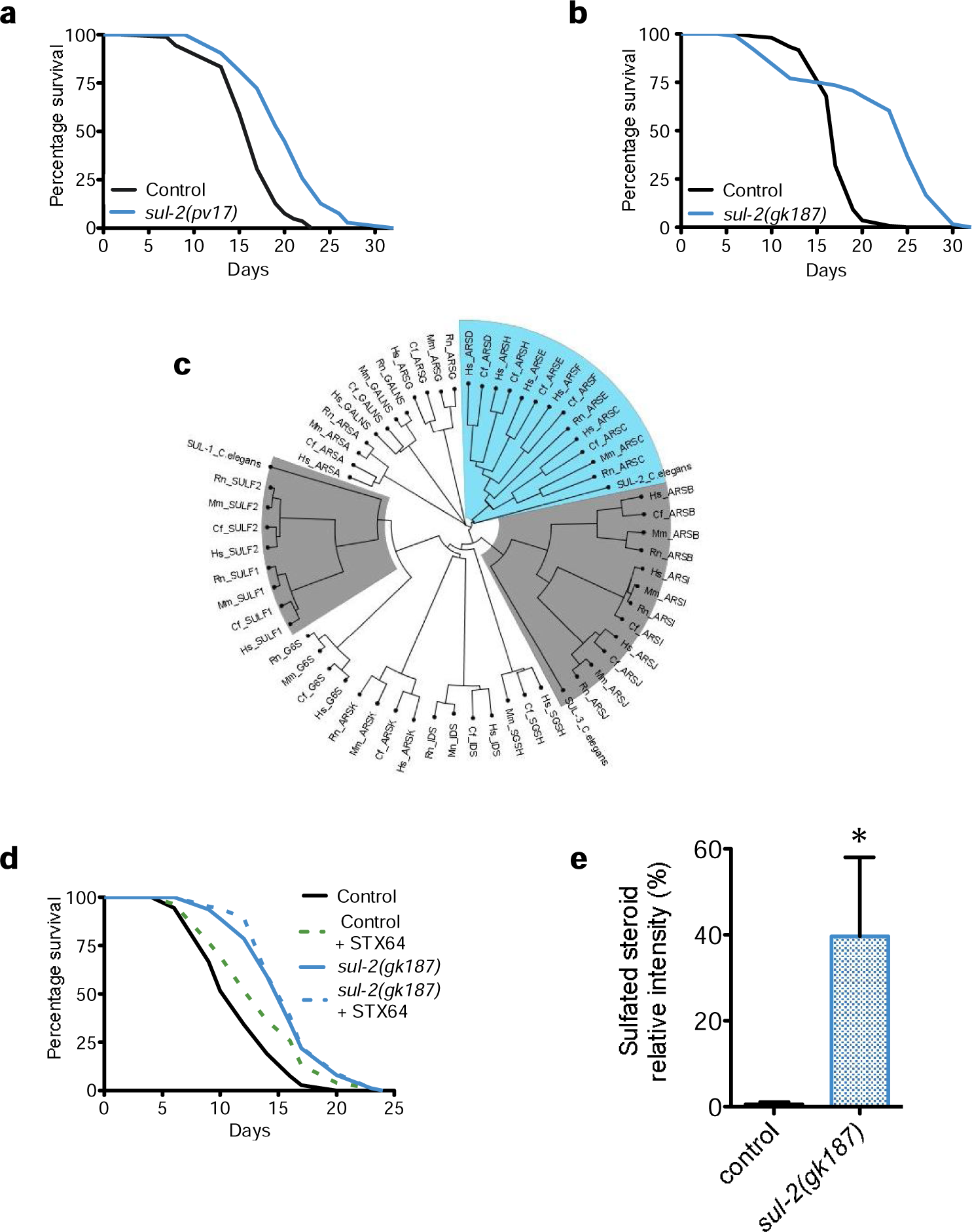
Reduction of activity of *sul-2* increases longevity and affects the levels of sulfated steroid hormones. **a**, *sul-2(pv17)* point mutation allele lives longer than wild type. **b**, The null allele *sul-2(gk187)* also increases lifespan. **c**, Phylogenetic tree of mammalian sulfatases and the three *C. elegans* sulfatases. Note that SUL-2 clusters with steroid sulfatases (type C) among others, in blue. Phylogenetic relation with other *C. elegans* sulfatases are indicated in grey. **d**, Inhibition of steroid hormones sulfatase by STX64 (1μg/ml) increases lifespan in wild type animals, but not in *sul-2(gk187)* background. Worms were cultivated in UV-killed *E. coli*. **e**, Deletion of *sul-2* generates an increase in the percentage of steroid hormones in the sulfated stage. Data from three independent assays are shown. One-tailed Mann-Whitney t-test. Statistics and additional longevity data are shown in Supplementary Table 1. Relative quantification of hormones from independent experiments are shown in Extended Data Table1.

Sulfatases are a large protein family involved in different biological processes and affect a wide range of substrates. The placement of curated *sul-2* in the sulfatases phylogenetic tree is uncertain, but when compared to mammalian sulfatases, *sul-2* clusters closer to the Arylsulfatases type H, F, E, D and the steroid sulfatase type C (Fig. 1c) that probably originated from a common ancestor gene^2^. We hypothesized that *sul-2* may exert its activity by modifying sulfated steroid hormones. Steroid hormone sulfatases are conserved proteins that participate, among other processes, stimulating proliferation in hormone-depending cancers^4^. Specific inhibitors for this type of enzymes have been generated, such as STX64^5^, which has been used to treat patients with hormone-dependending cancers^6^. We treated wild type animals with STX64 and observed an increase of longevity (Fig. 1d and Extended Data Fig. 3a). STX64 also phenocopies other *sul-2* mutant phenotypes (below and Extended Data Fig. 3b). STX64 does not further increase the longevity of *sul-2* deletion mutants, suggesting that the mechanism by which STX64 increases longevity is by inhibiting the sulfatase activity of SUL-2 (Fig. 1d).

In mammals, sulfated hormones have been long considered inactive forms that function mainly as reservoir and are activated by steroid sulfatases^4^, although a direct action of sulfated hormones in the nervous and reproductive system has been observed^7, 8^. Consequently, humans with STS deficiency show an increased blood levels of most sulfated steroids species^9^. We measured steroid hormones by a high-resolution HPLC-TOF-MS in *sul-2* mutants and found a higher proportion of sulfated hormones in this strain as compared to wild type worms (Fig. 1e and Extended Data Table 1). All these data suggest that SUL-2 can act as a steroid hormones sulfatase and regulate longevity through the alteration of the sulfated state of one or several steroid hormones.

To investigate if *sul-2* acts in a known longevity pathway, we performed genetic interaction studies with alleles that affect longevity. Mutations in the IGF receptor *daf-2* increase lifespan and this longevity requires the transcription factor DAF-16/FOXO^10^. *sul-2* mutations further extend the lifespan of *daf-2* reduction of function mutants (Fig. 2a and Extended Data Fig. 4a-b), suggesting that *sul-2* acts in a different pathway to regulate longevity. However, the increased longevity of *sul-2* mutants is mainly suppressed by DAF-16 loss-of-function (Fig. 2b). Longevity conferred by lack of germline also requires DAF-16^10, 11^, which translocates to the nuclei mainly in intestinal cells^8^. However, in insulin signalling mutants, DAF-16 localises to the nucleus of most cells^12^. In *sul-2* mutants, DAF-16 localises predominantly to intestinal nuclei (Extended Data Fig. 5a-b), suggesting a role for *sul-2* in germline-mediated longevity. Other essential factors for germline-mediated longevity, such as the intestinal ankyrin-repeat protein *KRI-1/KRIT-1* and the transcription elongation factor *TCER-1/TCERG1* are also required for a fully increased longevity of *sul-2* mutants, although low effect is observed with the nuclear hormone receptor NHR-80^13–15^ (Fig. 2c-e and Extended Data Fig. 4c-e). Moreover, loss of *sul-2* has no significant effect on longevity of the germline-less mutant *glp-1* or *mes-1* ^16^ (Fig. 2f and Extended Data Fig. 5c). All these data strongly indicate that *sul-2* mediates signalling from the gonad to regulate longevity. However, *sul-2* mutations do not affect fertility, reproductive age or gonad morphology (Fig. 2g and Extended Data Fig. 4d-g). Taken together, our findings indicate that *sul-2* affects a signal that regulates longevity to adjust lifespan to the reproductive status without affecting gonadal function.

**Figure 2.**
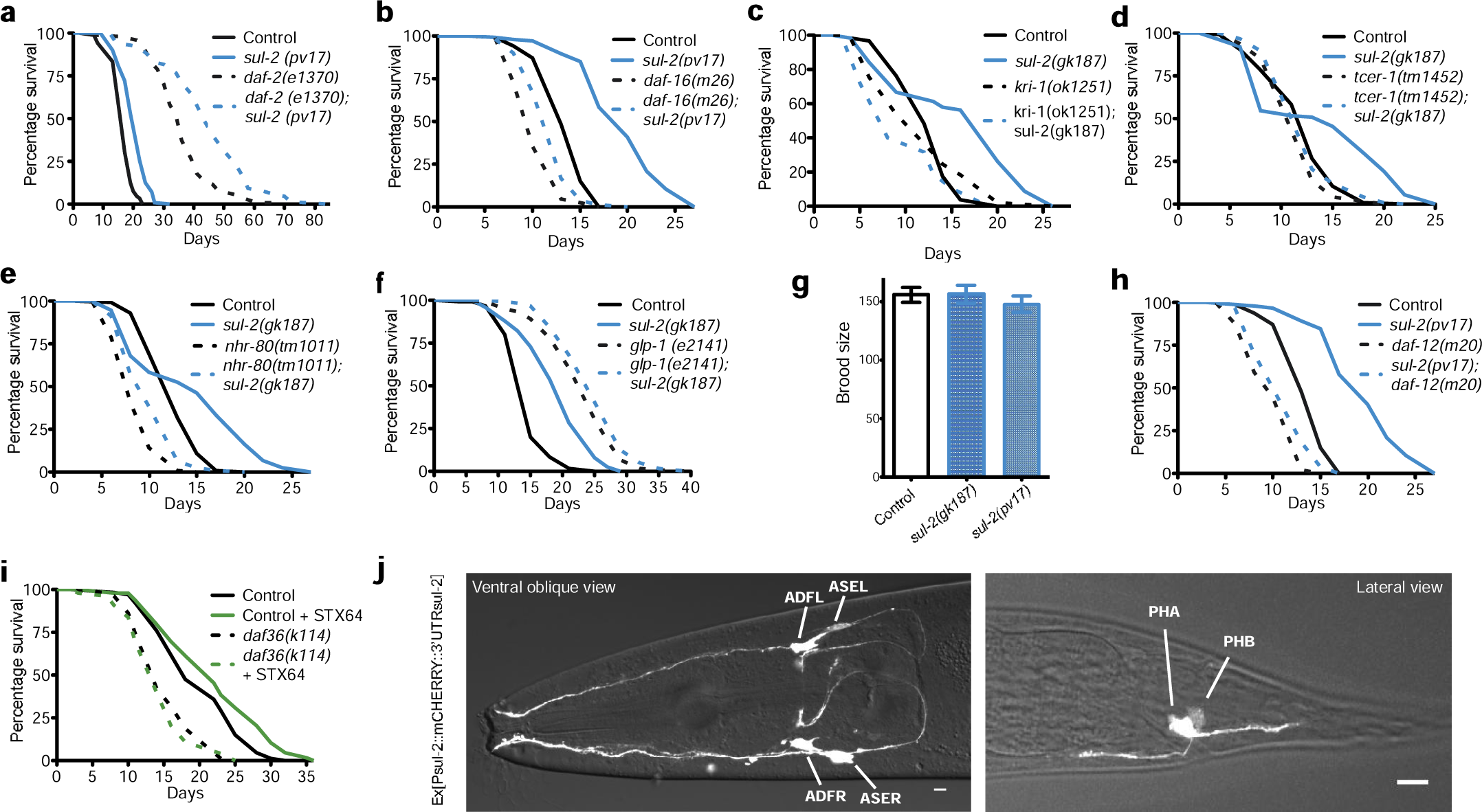
Genetic interactions and cellular location of *sul-2* expression. Genetic analysis show that *sul-2* mumimics longevity of animals without germline, but do not affect fertility. **a**, *sul-2* mutant enhances longevity in *daf-2(e1370)* background. **b**, *daf-16* transcription factor is required for *sul-2* longevity. **c, d, e**, Essential factors for germline-loss longevity *kri-1(ok1251)*, *tcer-1(tm1452)* and *nhr-80(tm1011)* are required for *sul-2* mutant longevity.**f**, *sul-2* deletion has not significant increase of longevity in *glp-1(e2141)* mutant background. **g**, Brood size of *sul-2* mutants are not significant different to wild type (25ºC). Mean±SEM; One-way ANOVA test. **h**, *daf-12* transcription factor is required for *sul-2* longevity. **i**, The Rieske-like oxygenase *daf-36* is necessary for the increase of longevity upon inhibition of steroid hormone sulfatase. **j**, *sul-2* is transcriptionally expressed in sensory neurons, mainly ADF and ASE in the head, and PHA and PHB in the tail. Scale bar 10 um. Statistics and additional longevity data are shown in Supplementary Table 1.

In germline-less animals the activation of the nuclear receptor DAF-12 by bile acid-like steroids called dafachronic acids (DAs)^17^ triggers an increase of longevity. We observed that *daf-12* is also needed for the increased longevity of *sul-2* mutants (Fig. 2h), indicating that *sul-2* inactivation causes the alteration of the sulfated steroid hormones pool, generating a signal upstream of DAF-12 that imitates the longevity of gonad depleted animals. DAF-36 converts cholesterol to 7-dehydrocholesterol in the first step of the biosynthetic pathway of △^7^-DA^18, 19^. Therefore, DAF-36 is also needed for the increased longevity of germline-less animals^20^. Similarly, DAF-36 is required for the longevity conferred by the steroid sulfatase inhibitor STX64 (Fig. 2i) placing the signal generated by sulfated steroid hormones upstream of the biosynthesis of DAs.

We have studied the anatomical location of *sul-2* expression in extrachromosomal array and single-copy insertion transgenic strains. We found that *sul-2* is expressed only in a few sensory neurons, mainly in the amphids ADF and ASE, and no detectable expression in germline were observed in any transgenic strains (Fig. 2j and Extended Data Fig. 6, 7a-b). ASE neurons are responsible for the attractive response of Na^+^ and Cl^−^, among others^21^. Defects in odour sensing affect longevity^22^. Therefore, we tested the ability of *sul-2* mutants to respond to Cl^−^ or Na^+^ and found no differences compared to wild type animals (Extended Data Fig. 7c-d). Furthermore, *sul-2* mutation increases the longevity of *daf-10(m79)*, a long-lived mutant defective in sensory cilia formation^22^ (Extended Data Fig. 7e). These results show that the longevity phenotype observed in *sul-2* mutants is not due to impaired functionality of sensory neurons.

Aging is considered the main risk factor for the onset of neurodegenerative disorders like Parkinson, Huntington, or Alzheimer. These disorders are caused by the progressive decline of proteostasis, which results in protein aggregation that compromises cellular function and finally causes cell death^23^. Germline-deficient *C. elegans* delay the symptoms derived from the proteotoxicity of ectopically expressed β-amyloid (βA)^24^. We tested if *sul-2* mutations or STX64 treatment improves the symptoms of *C. elegans* models for neurodegenerative diseases. In a *C. elegans* Parkinson disease model, α-synuclein expression in muscle cells cause age-dependent paralysis^25^, *sul-2* mutation or treatment with STX64 significantly improved mobility (Fig. 3a-b, Extended Data Fig. 8a-c and Extended Video 1-4). Loss of function of SUL-2 decreased the number of α-synuclein aggregates (Extended Data Fig. 8d-e), suggesting a better handling of protein aggregates in worms with reduced steroid sulfatase activity. To further assay the neuroprotective effect of reduced *sul-2* activity we tested a strain expressing α-synuclein in dopaminergic neurons. In this model, GFP-labelled dopaminergic neurons die due to α-synuclein toxicity^26^. Consistently, *sul-2* mutants showed increased neuron survival compared with control worms, indicating a neuroprotective action of reduced steroid-hormone sulfatase activity (Fig. 3c-d). In a Huntington neurodegenerative model expressing polyglutamine repeats fused to YFP, that aggregate in adult worms^27^, we found that both *sul-2* mutation and treatment with STX64 reduced the number of aggregates (Fig. 3e-f). We also tested an Alzheimer disease (AD) worm model where age-dependent immobility is caused by expression of βA protein in muscle cells^28^. Consistently, *sul-2* mutation and STX64 treatment delayed paralysis (Fig. 3g-h). All these data show that inhibition of *sul-2* protects against aging related proteotoxicity in the nematode.

**Figure 3.**
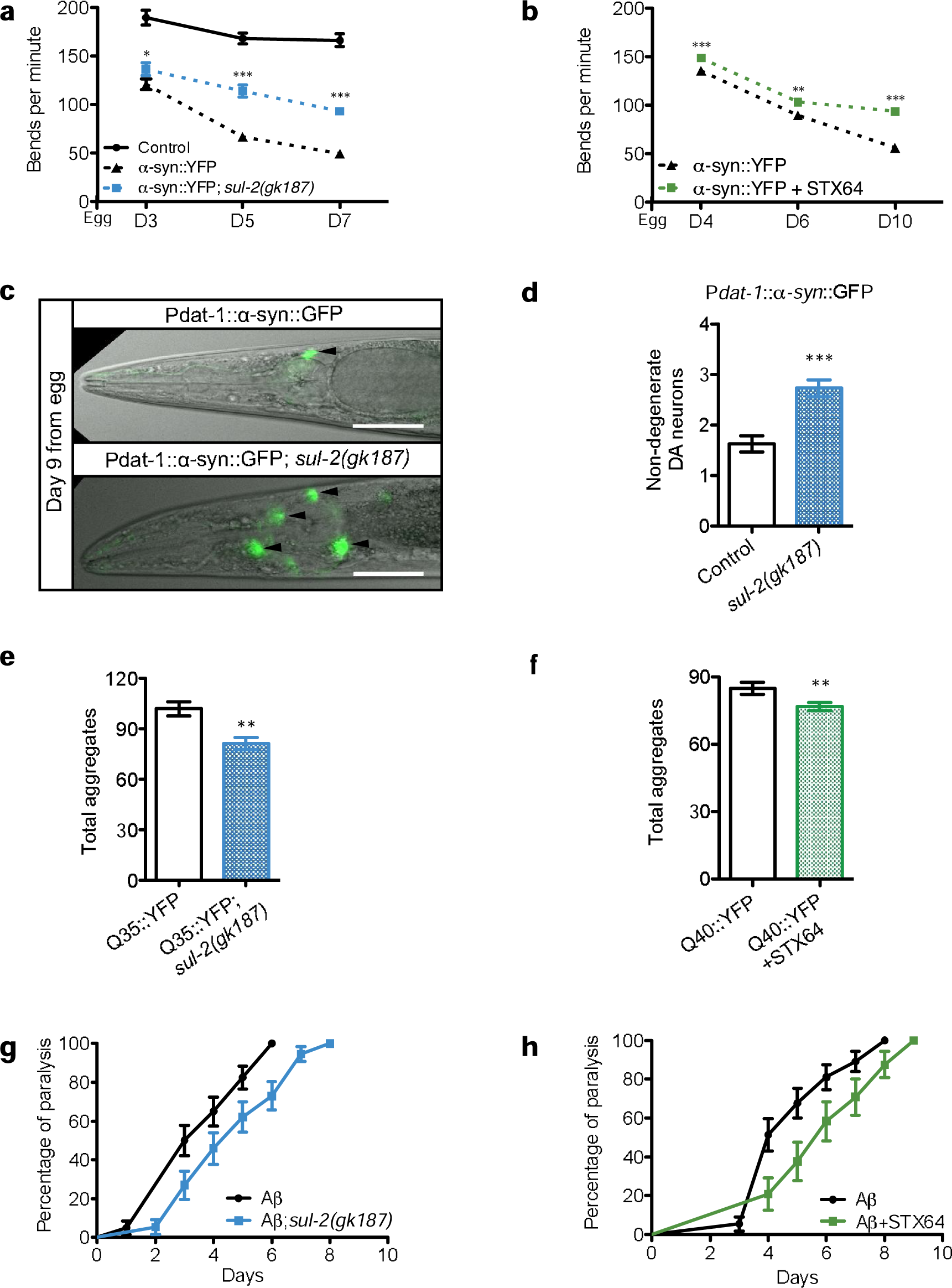
Reduction of steroid hormones sulfatase activity ameliorates the symptons of proteotoxicity models in C. elegans. **a**, Strains expresing a-synuclein in muscle cells reduce mobility with age at a slower rate in a *sul-2(gk187)* background (n ≥ 17) **b**, or under treatment with STX64 (n ≥ 9). **c, d**, Neurodegeneration of dopaminergic (DA) neuronsexpressing human α-synuclein is reduced in *sul-2* mutant background. A representative image of each condition and quantifications at day 9 old are shown. Data from two biological replicates are displayed, n=37. **e**, Q35 aggregates are reduced in *sul-2* deletion background (n ≥ 6), 8-day old worms. **f**, or by treatment with STX64 in Q40 background (n ≥ 12), 5-day old worms. **g**, Expression of hu¬man ß-amiloyd in muscles provokes paralysis with age that is amilorated in *sul-2* mutant back¬ground, **h**, or by STX64 treatment, log-rank (Mantel-Cox) test, for *sul-2* deletion p= 0.0027 and STX treatment p=0.007. Scale 50 μm. Additional biological replicate assays are shown in Ex¬tended Data Figure 8.

As STX64 ameliorated neurodegeneration in *C. elegans* models, we tested the effect of this drug on cognitive alterations provoked by intrahippocampal βA oligomers infusion, an acute AD mammalian model (Fig. 4a). Previously, it has been reported that local administration of DU-14, an inhibitior of steroid hormones sulfatase, could alleviate memory lost caused by intrahippocampal administration of βA oligomers in a mammalian model^29^. We observed that both, local and systemic STX64 treatments reverted cognitive deficiencies, measured by passive avoidance test, caused by intrahippocampal administration of βA oligomers. To evaluate the effect of STX64 oral treatment on amyloid pathology in a chronic AD mice model, we assessed the effect of 3-4 weeks of STX64 oral treatment on amyloid deposition in neocortex (the cerebral cortex and the hippocampus) of >15-month-old APP-PS1 mice (Fig. 4b). At this age corresponding to a late stage of amyloid deposition in the neocortex of the APP-PS1 model, the analysis of βA plaque density and size revealed a significant reduction, except for plaque size in hippocampus, in mice treated with STX64. Moreover, βA immunoreactive area is reduced in both tissues (Fig. 4c-e). Interesting, when we compared βA deposition in old (>15 month-old) APP-PS1 mice treated with STX64 with the normal temporal course of amyloid deposition in un-treated APP-PS1 mice, we observed that STX64 reduce βA deposition in APP-PS1 mice older than 15 month to that observed in APP-PS1 mice of 10-12 month of age (Fig. 4f). All these results indicate that STX64 treatment in APP-PS1 mice reduces βA deposition. We wondered if the histological improvement correlated with amelioration of cognitive behavioural deficit. For that we compared cognition capacity in APP-PS1 mice older than 15 month-old treated with vehicle or STX64 during 3-4 weeks. While vehicle-treated APP-PS1 mice showed a deficit in passive avoidance test (Fig. 4g), those mice treated with STX64 completely reverted cognitive deficiencies, reaching similar levels to <15 month-old wild type mice. All these results point out that the alterations in βA metabolism provoked by STX64 reduce the cognitive behaviour deficiencies induced by βA accumulation in acute and chronic AD mice models, suggesting a potential for STX64 as a pharmacological therapy against neurodegenerative diseases.

**Figure 4.**
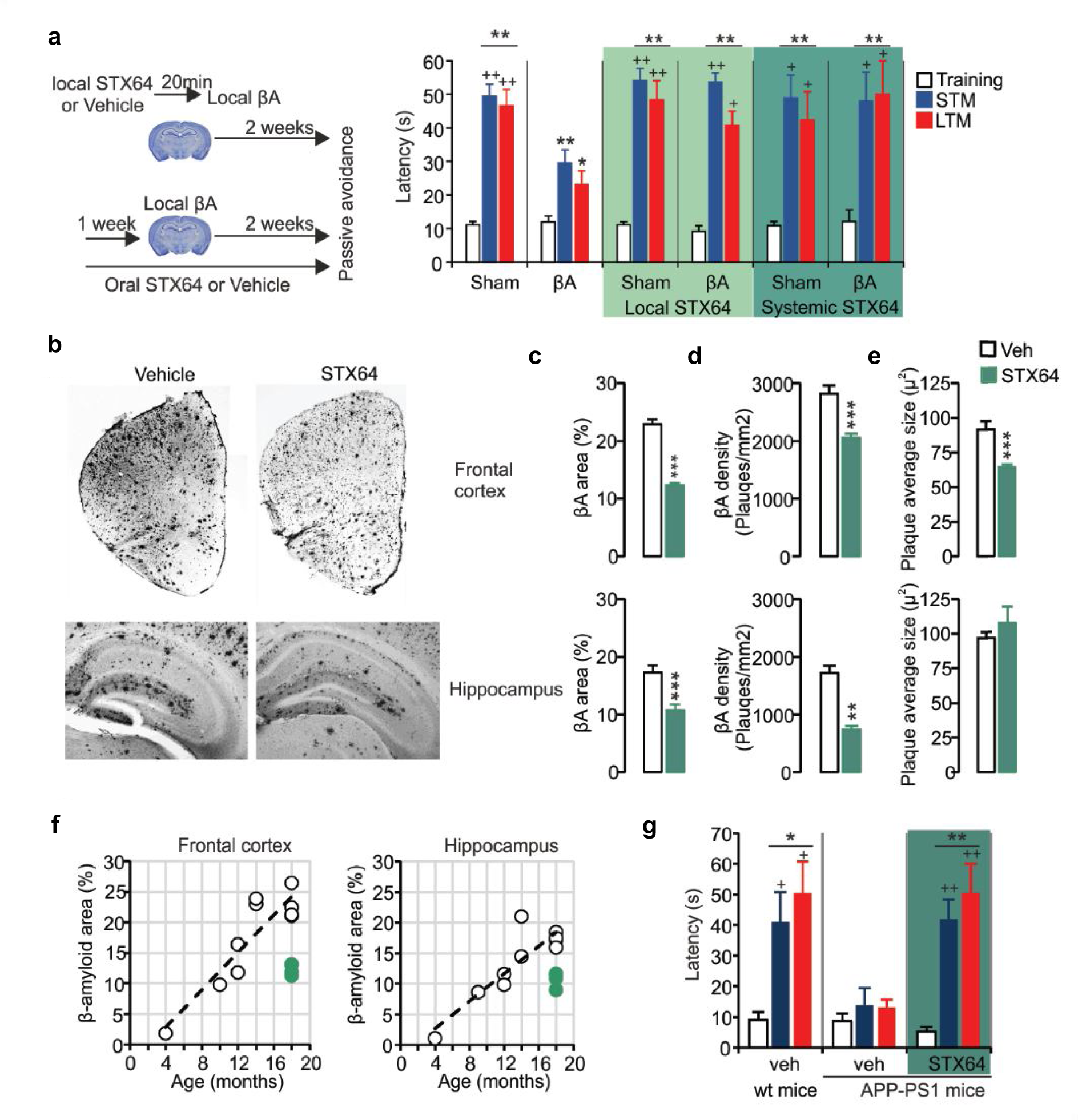
STX64 treatment alleviated ß-amyloid (ßA) deposition and cognition deficiency in Alzheimer disease mice models. **a**, Effect of intrahippocampal (1pM) and oral administration (5 pg/ml in drink water) of STX64 in the passive avoidance test in wild type mice injected with ßA oligomers in the hippocampus. The number of mice in each group were >5. **b**, Representative ßA-immunoreactive images assessed in the frontal cortex and the hippocampus of APP-PS1 mice older than 15 months of age after 3-4 weeks of vehicle or STX64 intake. **c**, Quantification of percentage of ßA area, **d**, deposition density and **e**, average plaque size in the frontal cortex and the hippocampus of >15 months-old APP-PS1 mice after 3-4 weeks of oral administration with STX64 or vehicle. n=4 mice per groups. **f**, Temporal course of ßA deposition in APP-PS1 mice and the effect of 3-4 weeks STX64 oral treatment on ßA area in the frontal cortex and the hippocampus. Green dots represent STX64 treated animals, The number of microphotographs quantified were more than 3 in each mice tested. **g**, Effect of oral administration with STX64 in more than 15-month-old APP-PS1 mice, and comparation with APP-PS1 and wild type mice older than 15 months in the passive avoidance test. The number of mice in each group were >5. In histological analysis, * represents significant differences between vehicle-and STX64 administered APP-PS1 mice. In behavioural test, * represents significant differences between the short-term and long-term memory sessions (STM and LTM, respectively) with respect the training session in the same experimental group; and, ^+^ represents significant differences between the STM and LTM sessions between each experimental group and ßA group. A symbol, p<0.05; two symbols, p<0.01; and three symbols, p<0.001.

Germline stem cells inhibit, through an unknown mechanism, a pro-longevity signal generated from the somatic gonads. Our data suggests that steroid hormones are implicated in this process. Enzymes involved in the sulfate modification of steroid hormones (sulfatase SUL-2 and sulfotranferase SSU-1) are expressed in sensory neurons^30, 31^. This implies that alteration of hormones sulfate state may act in the integration of environmental cues, such as nutrient availability, with the reproductive status, which are two-linked process^32, 33^. We have shown that alteration of sulfated state impinge in longevity and in aging related diseases. We cannot sort out whether it is the lack of non-sulfated hormones or the presence of sulfated hormones what has a beneficial effect (See model in Extended Data Fig. 9). Regulation of steroid hormones sulfation is a conserved process. In humans dehydroepiandrosterone sulfate (DHEAS) declines with age and has been used as a marker of aging, raising speculations of a causative effect on sarcopenia, poor cognitive function and other aging associated diseases^4^. Our data show that STX64 treatment extends lifespan in *C. elegans* and protects against aging-associated proteotoxicity in nematodes and mice. As a result of this observation, inhibitors of steroid hormones sulfatase, such as STX64, could be pharmacological compounds of interest to be reallocated for treating aging associated diseases.

## Acknowledgements

We thank the CGC, A. Miranda-Vizuete and J. Alcedo for providing strains. Y. Kohara and P. Askjaer for DNA clones. P. Askjaer and A. Garzón for critical review of the manuscript. A. P. Pulido for bioinformatic advices. V. Carranco, for his excellent technical assistance during whole project and also A. López, A. Cano, V. Rubio, S. Romero and E. Gara for their technical support. K. Garcia for microscopy assistance. This work was supported by the Junta de Andalucía Project P07-CVI-02697 and the European Research Council (ERC-2011-StG-281691).

## Author Contributions

M.M.P.J. and M.J.M. conceived and designed the study, and the *C. elegans* experiments. M.M.P.J. performed the *C. elegans* experiments. M.M.P.J., P.S and A.V. did the neurodegeneration assays in *C. elegans*. J.M.M. isolated *pv17* allele. A.M.C., S.E.G., and I.S.P. performed mice assays. A.M.C conceived the mice and J.J.R. performed hormone identification assays. M.M.P.J., A.M.C., and M.J.M performed formal analyses and data presentation. M.M.P.J., M.A.S., A.M.C., and M.J.M. interpreted results and wrote the manuscript.

## Methods

### Strains

All the strains were grown under standard conditions^34^. **N2:** wild type Bristol isolate, **GM11:** *daf-16(m26)I;fer-15(b26)II*, **\*GM88:** *daf-16(m26)I;sul-2(pv17)V*, **\*GM369:** *kri-1(ok1251)I*, **\*GM354:** *kri-1(ok1251)I;sul-2(gk187)V*, **\*GM357:** *kri-1(ok1251)I;sul-2(pv17)V*, **\*GM363:** *daf-16(mu86)I; muIs109[Pdaf-16::GFP::DAF-16 cDNA+Podr-1::RFP]X*, **\*GM365:** *daf-16(mu86)I;sul-2(pv17)V;muIs109*, **\*GM366:** *daf-16(mu86)I;sul-2(gk187)V;muIs109*, **AM140:** *rmIs132[Punc-54::Q35::YFP]I*, **\*GM387:** *rmls132;sul-2(gk187)V*, **AM141:** *rmIs13[Punc-54::Q40::YFP]*, **EG4322:** *ttTi5605II;unc-119(ed9)III*, **\*GM410:** *pvIs8[Psul2::mCHERRY::3’UTR sul-2]II*, **CL2006:** *dvIs2[pCL12*(P*unc-54::Aß1-42::3’unc-54)+pRF4(rol-6(su1006)]II*, **\*GM392:***dvIs2;sul-2(gk187)V*, **DH26:***fer-15(b26)II, **GM6:** fer-15(b26)II; daf-2(e1370)III*, **GM270:** *fer-15(b26)II; daf-2(m577)III*, \***GM125:** *fer-15(b26)II;sul-2(pv17)V*, **\*GM325:** *fer-15(b26)II;sul-2(gk187)V*, **\*GM293:** *fer-15(b26)II;daf-2(m577)III;sul-2(pv17)V*, **CF2167:***tcer-1(tm1452)II*, **\*GM348:***tcer-1(tm1452)II;sul-2(gk187)V*, **\*GM353:***tcer-1(tm1452)II;sul-2(pv17)V*, **GM63:***daf-2(e1370)III*, **DR1942:** *daf-2(e979)III*, \***GM134:** *daf-2(e1370)III;sul-2(pv17)V*, **\*GM291:***daf-2(e1370)III;sul-2(gk187)V*, **\*GM215:** *daf-2(m577)III;sul-2(pv17)V*, **\*GM229:** *daf-2(e979)III;sul-2(pv17)V*, \***GM407:***glp-1(e2141)III*, \***GM408**: *glp-1(e2141)III;sul-2(gk187)V*, **NL5901:** *pkIs2386[unc-synuclein::YFP + unc-119(+)]III*, **\*GM379:** *pkIs2386;sul-2(gk187)V*, **BX165:** *nhr-80 (tm1011)III*, **\*GM360:** *nhr-80(tm1011)III;sul-2(gk187)V*, **\*GM358:** *nhr-80(tm1011)III;sul-2(pv17)V*, **PR678:** *tax-4(p678)III*, **NY2067:** *ynIs67[Pflp-6::GFP]III;him-5(e1490)V*, **\*GM314:** *ynIs67[Pflp-6::GFP]III;pvEx3*, **QZ65:** *daf-10(m79)IV*, **\*GM351:** *daf-10(m79)IV;sul-2(gk187)V*, \***GM126:** *sul-2(pv17)V*, **\*GM371:** *sul-2(gk187)V*, **\*GM272:** *sul-2(pv17)V;daf-12(m20)X*, **\*GM364:** *sul-2(gk187)V;mes-1(bn7)X*, **DR20:** *daf-12(m20)X*, **\*GM359:** *mes-1(bn7)X*, **VZ155:** *oyIs26[Pops-1::GFP]X*, **\*GM318:***oyIs26[Pops-1::GFP]X;pvEx3*, **\*GM288:***pvEx1[Psul-2::mCHERRY::3’UTR sul-2 + pGK10(sca-1::GFP)]***\*GM303:**N2;*pvEx3[Psul-2::mCHERRY::3’UTR sul-2]*, **GR1366:** *mgIs42 [tph-1::GFP + pRF4(rol-6(su1006))]*, **\*GM320:** *mgIs42;pvEx3*, **UA44:** *baln11 [Pdat-Syn::unc-54 3’-UTR; Pdat-1::GFP]*, **\*GM391:** *fer-15(b26)II; sul-2(gk187)V; baln11.* * Indicate the strains backcrossed with our N2 or generated during the study.

### Mutant isolation

In order to obtain the *sul-2(pv17)* mutant strain we performed a mutagenesis protocol as described in Muñoz and Riddle (2002)^1^. Briefly: The temperature-sensitive fertilization-defective mutant *fer-15(b26*ts), was treated with ethyl methanesulfonate (EMS) as described^35^. Sets of 30 mutagenized L4 larvae were incubated for 8 days at 15ºC in tubes containing 6 ml S medium with *E. coli^36^*. F2 eggs were purified by alkaline hypochlorite treatment to obtain synchronous L1 larvae^37^. Approximately 10.000 synchronous F2 or F3 L1 larvae were harvested from each tube and submitted to thermal stress at 30ºC for 7 days on agar plates spread with OP50. To ensure independence of the mutants, only one survivor per tube was saved after confirming L1 thermotolerance at 30ºC. One of those mutants isolated was *sul-2(pv17)* allele.

### *pv17* identification

To map the mutation *sul-2(pv17)*, we used a snp-SNPs based method with the Hawaiian-type strain as described in Wicks et al. (2001)^38^, combined with classical genetic markers. Using this approach, we found that *sul-2(pv17)* is located between SNP CE5-168 at 1.00 cM and SNP pkP5062 at 1.11 cM on chromosome V, equivalent to a physical distance of 205 Kb. Then, using transgenic lines we tried to complement the mutation with genes of this region, found that the wild type *sul-2* gene complemented the mutant *sul-2(pv17)*. Sequencing of the wild type and *sul-2(pv17)* allele identified a mutation in the gene which consists in a missense mutation that change a glycine to an aspartic acid residue at the position 46. An EST clone of the *sul-2* gene (yk387h10) was sequenced. The protein obtained differs from the one published in wormbase (http://www.wormbase.org) (see Extended Data Figure 2).

### Lifespan assays

Strains were synchronized by hypochlorite treatment of gravid adults, grown up during two generations at 16 ºC. F2 L4 animals were shifted to 25ºC, unless otherwise indicated (t=0). During the first week animals were transferred to a fresh plate every two days, further they were transferred at least once per week and scored every two days until death. Animals lost or dead by non-physiological causes were censured. GraphPad Prism 5 (Version 5.0a) was used to analyse the data. Survival curves were generated using the product-limit method of Kaplan and Meier. The log-rank (Mantel– Cox) test was used to evaluate differences in survival and p-values lower than 0.01 were considered significant. For STX64 (Sigma-Aldrich, Ref. S1950) lifespan assays starting at L4 stage, control and treated plates were prepared fresh every two days and animals were transferred. STX64 was disolved in DMSO. Controls of all lifespan assays were *fer-15(b26)* mutant background, unless otherwise indicated, to avoid progeny production. *fer-15(b26)* mutation does not have effect on life span^39^. For lifespan details see Supplementary Table 1.

### Phylogenetic analysis

Phylogenetic analysis was based on the *C. elegans* and mammal sequences described in Sardiello et al. 2005^2^ and multiple sequence alignments were performed by MAFFT^40^, which is available at http://www.ebi.ac.uk/Tools/mafft/. e Default parameters were used. Phylogenetic tree was visualized using the program FigTree v.1.4.3 (http://tree.bio.ed.ac.uk/software/figtree/).

### Steroid purification, high-performance liquid chromatography (HPLC-MS) and data analysis

~100 000 worms were collected in a polypropylene Falcon tube. The worms were homogenized in M9 buffer/methanol (2/3, v/v). For steroid quantification 11-deoxycortisol-2,2,4,6,6-d5 (710784; for unconjugated steroids), sodium pregnenolone-17A, 21,21,21-d4 sulfate (721301; for sulfated steroids) and diethylhexyl phthalate-d4 (DEHP-d4) were introduced into the extract. The sample was then sonicated in an ultrasonic bath for 5 min, left overnight at room temperature and centrifugated at 3000 g for 5 min. The organic phase was collected and the rest of the extract residue washed again with 5 ml of methanol containing 1% acetic acid and centrifuged. The two organic phases were pooled and evaporated to dryness at 40 °C under a gentle stream of nitrogen, taken up in 1 ml of acetonitrile. The sample was purified by vortex-mixing for 1 min and centrifuged at 10.000 × g for 10 min. The supernatant was transferred to another 1.5 mL Eppendorf tube and evaporated to dryness under nitrogen atmosphere at 40 °C. –water (5:95, v/v). A –MS/MS system for analysis.

The liquid chromatograph in the HPLC/ESI-TOF-MS system was Dionex Ultimate3000RS U-HPLC (Thermo Fisher Scientific, Waltham, MA, USA). Chromatographic separation was performed as follows. The eluent components were 0.1% (v/v) formic acid in water (A), and 0.1% (v/v) formic acid in methanol (B). The proportion of B was increased from 50% to 75% in 12 min and held 25 min, then increase to 95 % in 5 min and held for 5 min. Initial conditions were reached in 5 min and the equilibrium time was 2 min. The injection volume was 30 µL and the flow rate was 1 mL/min. A stainless steel column (20 × 0.46 cm i.d.), packed with 3µm C18 Spherisorb ODS-2 (Teknokroma, Barcelona, Spain) was used. A split post-column of 0.4 mL/min was introduced directly on the mass spectrometer electrospray ion source. Mass spectrometry was performed using a micrOTOF-QII High Resolution Time-of-Flight mass spectrometer (UHR-TOF) with q-TOF geometry (Bruker Daltonics, Bremen, Germany) equipped with an electrospray ionization (ESI) interface. All data were used to perform multitarget-screening using TargetAnalysis^TM^ 1.2 software (Bruker Daltonics, Bremen, Germany). Collision energy was estimated dynamically based on appropriate values for the mass and stepped across a +/-10% magnitude range to ensure good quality fragmentation spectra. The instrument control was performed using Compass 1.3 for micrOTOF-Q II+Focus Option Version 3.0.

The *in-house* mass database created *ex professo* comprises monoisotopic masses, elemental composition and, optionally, retention time and characteristic fragment ions if known, for steroids and their derivatives compounds (Supplementary Table 2). Data evaluation was performed with Bruker Daltonics DataAnalysis 4.1. From the HPLC/TOF-MS acquisition data, an automated peak detection on the EICs expected for the [M+H]^+^ ions of each compound in the database was performed with Bruker Daltonics TargetAnalysis^TM^ 1.2 software. The software performed the identification automatically according to mass accuracy and in combination with the isotopic pattern in the SigmaFit^TM^ algorithm. This algorithm provides a numerical comparison of theoretical and measured isotopic patterns and can be utilized as an identification tool in addition to accurate mass determination. The calculation of SigmaFit values includes generation of the theoretical isotope pattern for the assumed protonated molecule and calculation of a match factor based on the deviations of the signal intensities. Only those hits with mass accuracy and SigmaFit values within the tolerance limits, which were set at 5 ppm and 50, respectively, are included in the final report list that was carried out using a Microsoft EXCEL-based script. The interpretation of the MS spectra was performed using the SmartFormula3D^TM^ module included in the DataAnalysis software. Based on expected chemistry, carbon elements, hydrogen, oxygen, nitrogen, bromine and iodine were permitted. Sodium and potassium were also included for the calculation of adduct masses. The number of nitrogen atoms was limited to an upper threshold of ten. The number of rings plus double bonds was checked to be chemically meaningful (between 0 and 50). For each steroid compound detected in the sample, the module shows the original MS and MS-MS data as peak lists. From all possible formulae for the precursor ion, only one should fit with the elemental composition expected for the [M+H]^+^ ion and satisfy thresholds for mass accuracy and SigmaFit values. Once the correct formula is selected, the module displays the formulae and neutral losses in the MS-MS spectrum fitting to the boundary conditions for the precursor ion, and they should be consistent with the MS-MS data peak list. The SmartFormula3D checks the consistency highlighting the monoisotopic peaks with formula suggestion and the related isotopic peaks. Based on this combined data evaluation, fragmentation pattern for each steroid hormone can be generated to support its identification in the sample.

### Brood size

Animals were grown at 16ºC. For the assays, 10 individual L4 animals of each strain were incubated at 20ºC or 25ºC, and monitored during all reproductive period. Animals were transferred to a fresh plate every single day and numbers of eggs were counted until animals did not lay more eggs.

### *sul-2* expression

To create the DNA construction for the transgenic strains, we amplified 1,7 Kb upstream of *sul-2* start codon (*5’AAAGATTTTTAACTGCCGTTTTTC3’-5’GATCTGAAAGATTATGAATGAAATCAA3’*) named P*sul-2* fragment and 0.3 Kb downstream of *sul-2* stop codon (*5’AT**GGATCC**GAATTTCTAAAATTC3’* -*5’TA**GGATCC**ATTTTGTAAATTAGCAC3’*) named 3’UTR *sul-2*, from wild type DNA. Both regulatory elements were transitory cloned into pGEM-T Easy Vector (Promega), next inserted into the pCR™-Blunt II-TOPO™ Vector (Invitrogen) containing mCHERRY (plasmid generated and kindly provided from P. Askjaer’s group, used previously to generate pBN1^*41*^. P*sul-2* introduced upstream of mCHERRY into *ApaI/PstI and* 3’UTR *sul-2* downstream in BamHI site (pPV5). Extrachromosomal strains were generated by microparticle bombardment^42^ of N2 with pPV5 together with pGK10 as cotransformation marker (GM288: *pvEx1*) and by microinjection of plasmid pPV5 into N2 (GM303: pvEx3). We observed the same expression pattern in all transgenic strains. For identification of *sul-2* neurons adults animals staining with FiTC or crossed with specific neuronal marker strains were imaging in Confocal Microscope Leica SP2-AOBS. To generated the integrated strain GM410 we amplified the whole casette P*sul-2*::mCHERRY::3’UTR *sul2* from pPV5 and inserted it, first in pGEM-T Easy Vector, next into *SbfI/SphI* of *pBN8* recombination plasmid modified for Mos1-mediated single-copy insertion (MosSCI) into chromosome II^41^ (pPV7). The transgenic animals were generated by microinjection of pPV7 with pBN40, pBN41 and pBN42 as green co-markers and pJL43.1 for transposase^43, 44^. Integrated animals were selected as wild type locomotion and non co-markers expression individuals. Pictures of adult animals were taken in Fluorescence Mi-croscopy Zeiss Axio Imager M2.

### Thrashing assay

Nematodes were synchronized to L4 - young adult at 20°C (3-day old worm) and thrashing were assayed at the indicated days. For each thrashing assay, a nematode was placed in a drop of M9 buffer, let 30 seconds for accommodation and later counted the number of bends during 1 minute. We considered a bend when the head of the nematode crosses the longitudinal axis. Number of animals assayed per day and condition are indicated in figure legends.

### Dopaminergic neurodegeneration induced by α-synuclein

Synchronized nematodes were grown up to day 9 (6 days of adulthood) and imaged in Fluorescence Microscopy Zeiss Axio Imager M2. For quantification of non-degenerate dopaminergic (DA) neurons we considered as normal neuron those where cell body and neurites are present, criteria based on Harrington et al. (2011)^45^.

### Quantification of aggregates

Animals were cultivated at 20ºC and imaged in Leica Fluo III stereoscope or Fluorescence Microscopy Zeiss Axio Imager M2. For polyQ strains and conditions, manual quantification of total aggregates of the whole animal have been done. To quantify α-synuclein aggregates total number of fluorescent aggregates between the two pharyngeal bulbs were counted.

### Paralysis assay

Paralysis was monitored during adulthooh (t0= first day of adult). Paralysis was considered when nematodes did not move after stimulation with a platinum pick. For each test condition, 50 nematodes were analyzed. Nematodes were synchronized and assayed at 20°C.

### Mice strains and conditions

The male Swiss (CD1) and APP-PS1^46^ mice used in this study were purchased from an authorized provider (University of Seville, Spain) and they were habituated to standard animals housing conditions for 2-3 weeks (a 12 h light/dark cycle, temperature and humidity). Behavioural studies were performed with 8 weeks-of-age Swiss mice, and >15 month-old in APP-PS1 mice in C57Black background. For histological studies, male APP-PS1 mice from 2 to >15 month-old were used. All experiments were performed in accordance with European Union guidelines (2010/63/EU) and with Spanish regulations for the use of laboratory animals in chronic experiments (RD 53/2013 on the care of experimental animals: BOE 08/02/2013), and the approval of the University Pablo de Olavide animal care committees was obtained prior to performing this study.

### Mice local drug infusion

Mice were anesthetized with 4% chloral hydrate (10 µL/kg of body weight, i.p.) and when fully anesthetized, they were situated in a stereotactic frame. In order to injure the hippocampus, 05μl of 5μM solution of β-amyloid (βA) oligomers were injected bilaterally into the dorsal hippocampus of the mice at the following stereotactic coordinates: AP = −2.2 mm, ML = ±1.5 mm, V = −1.5 mm from the Bregma. The mice were then allowed to recover for at least 15 days. Those mice that received also STX64, 20 minutes before βA oligomers were administrated with 0.5 μl of STX64 (1 mg/ml) were infused in the same rostral hippocampus coordinates. STX64 and βA oligomers were delivered at a rate of 0.2μl/min through an injection syringe(Hamilton), and left in place for 2.5 min following infusion.

### Mice Oral STX administration

STX64 was dissolved in drinking water at 0.005mg/ml. Mice were exposed to STX solution during 3-4 weeks and the water intake were registered every day during the treatment. The estimated STX64 doses were between 1-2 mg per Kg of mice and day.

### Step-through passive avoidance (PA) test

Mice have an innate preference for dark and enclosed environments. During the habituation phase, mice were handled and allowed to move freely for 1 minute in a chamber (47×18×26 cm, manufactured by Ugo Basile) that is divided symmetrically into one light and one dark compartment (each measuring 28.5 cm×18 cm×26 cm). During the training phase, mice were briefly confined to the light compartment and then 30 seconds later, the door separating the dark-light compartments was opened. Once mice entered the dark compartment, the door was closed automatically and the mice received an electrical stimulation (0.5mA, 5s and 0.3mA, 5s for Swiss and C57Black mice, respectively) delivered through the metal floor. In the retention tests performed at the times indicated, mice that recalled the electrical shock experience when re-placed in the light compartment avoided, or at least took longer, to enter the dark compartment. Thus, the latency to enter into the dark compartment (escape latency) is a measure of information learning or memory retention depending on how long after the training session the test was carried out. Escape latency (s) the training, short- and long-term memory (STM and LTM, respectively) sessions are represented.

### Immunohistochemistry and histological analysis

For immunohistochemistry, an antibody against βA (1:3000, Clone BAM-10, Sigma Aldrich Ref: A3981) was used. Antibody staining was visualized with H2O2 and diaminobenzidine, and enhanced with niquel. To minimize variability, at least 5 sections per mice were analyzed under a bright-field DMRB RFY HC microscope (Leica). In each section, the percentage area occupied by βA, the density and the average size of βA accumulations were quantified using Image-J software (downloaded as a free software package from the public domain: http://rsb.info.nih.gov/ij/download.html).

---

**Extended Data Figure 1.**
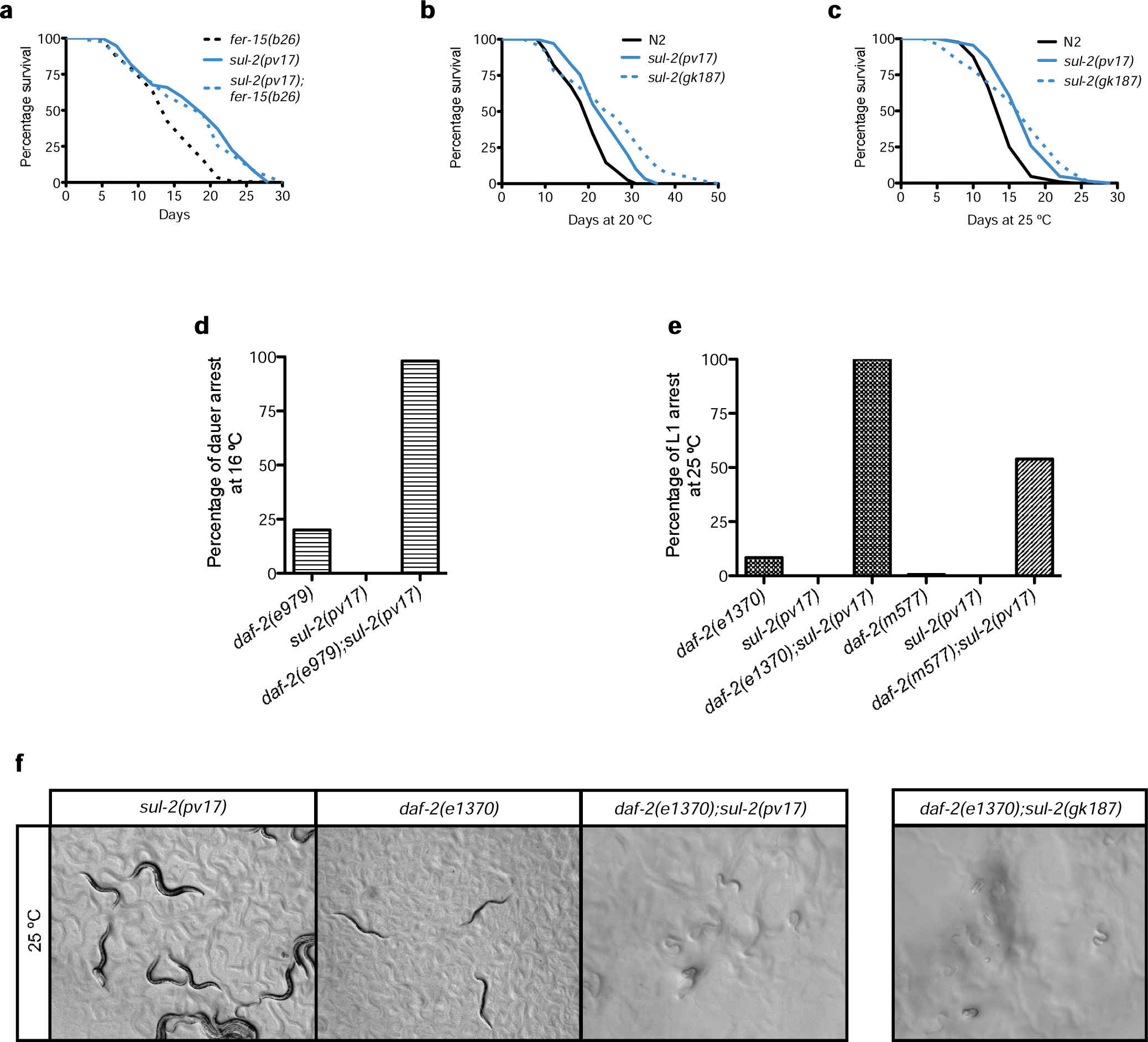
Longevity assays of *sul-2* and genetic interactions with *daf-2*. *sul-2* mutants do not show visible phenotypes, but are long-lived and enhance developmental phenotypes of *daf-2* mutants. **a**, *sul-2is* long-lived and *fer-15(b26)* does not affect their lifespan. **b**, *sul-2* mutants are long-lived at 20 °C. **c**, and at 25 °C. **d**, A small percentage of *daf-2(e979)* animals arrest development in dauer larvae at 16 °C, *sul-2(pv17)* mutant does not show any larval arrest, but enhances dauer arrest of *daf-2(e979).* **e**, Most *daf-2 (e1370)* animals arrest in dauer stage when develop at 25 °C, but small percentage arrest in L1 stage. In this condition, all animals from *daf-2(e1370); sul-2(pv17)* double mutants arrest at L1 stage. Similarly, more than 50% of animals arrest in L1 stage in *daf-2(m577); sul-2(pv17)* background, while none of the single mutants show this phenotype. **f**, Example of *sul-2(pv17)* larvae, dauer larval arrest of *daf-2(e1370)* or L1 arrest of *daf-2(e1370); sul-2(pv17)* and *daf-2(e1370); sul-2(gk187)* at 25 °C. Animals were grown up to L4 at 16 °C, then shifted at 25 °C, progeny were scored or imaged after 72 hours. Photographs were taken in a Leica scope. Statistics and additional longevity curves are shown in Supplementary Table 1.

**Extended Data Figure 2.**
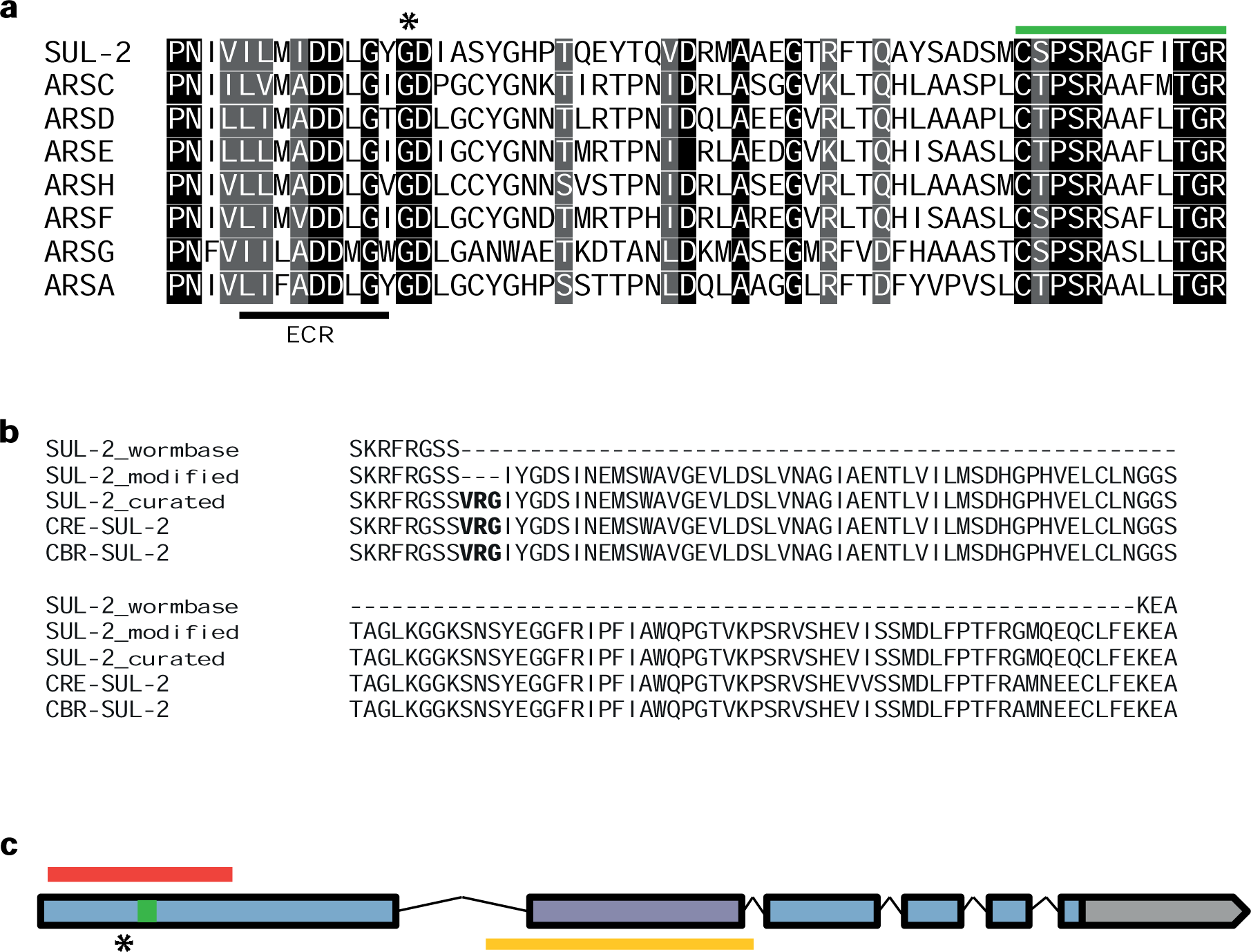
Identification of *pv17* allele and curated sequence of SUL-2. **a**, The *pv17* allele is a missense mutation that changes the glycine indicated with asterisk to an aspartic acid residue. The mutation is located close to an evolutionary constrained region (ECR)^2^, indicated with the bar at the bottom and also near to the catalytic core of sulfatases (in green). b, The DNA sequence we identified in the wild type *sul-2* differs to the one published in wormbase (SUL-2_wormbase) and identified as ortho-logue to ARSA (http://www.wormbase.org), this sequence misses 459 bp, also described recently by Li et al. (2015) ^3^ and predicted as part of an new exon [wormbase_170818 gw3] based on RNAseq data (SUL-2_modified). The protein sequence obtained by cDNA sequencing of yk387h10 clone (SUL-2_cu-rated) differs to the one predicted in three aminoacids (in bold). Notice that the three aminoacids identified in this sequence are present also in other species (CRE: *C. remanei*, CBR: *C. brigssae).* c, Exon intron composition of wild type *sul-2* and region deleted in the *gk187* allele (in red). The mutant lesion is available at wormbase(http://www.worm-base.org/species/c_elegans/variation/WBVar00145594#02-45-3). This allele deletes the sequence that encodes to the catalytic core of sulfatases (in green) and generates a frame shift, conserving only the four first aminoacids of the original sequence; therefore we considere *gk187* a null allele. The *pv17* allele location is indicated with asterisk, the new DNA fragment indentified in yellow and the new exon in purple.

**Extended Data Figure 3.**
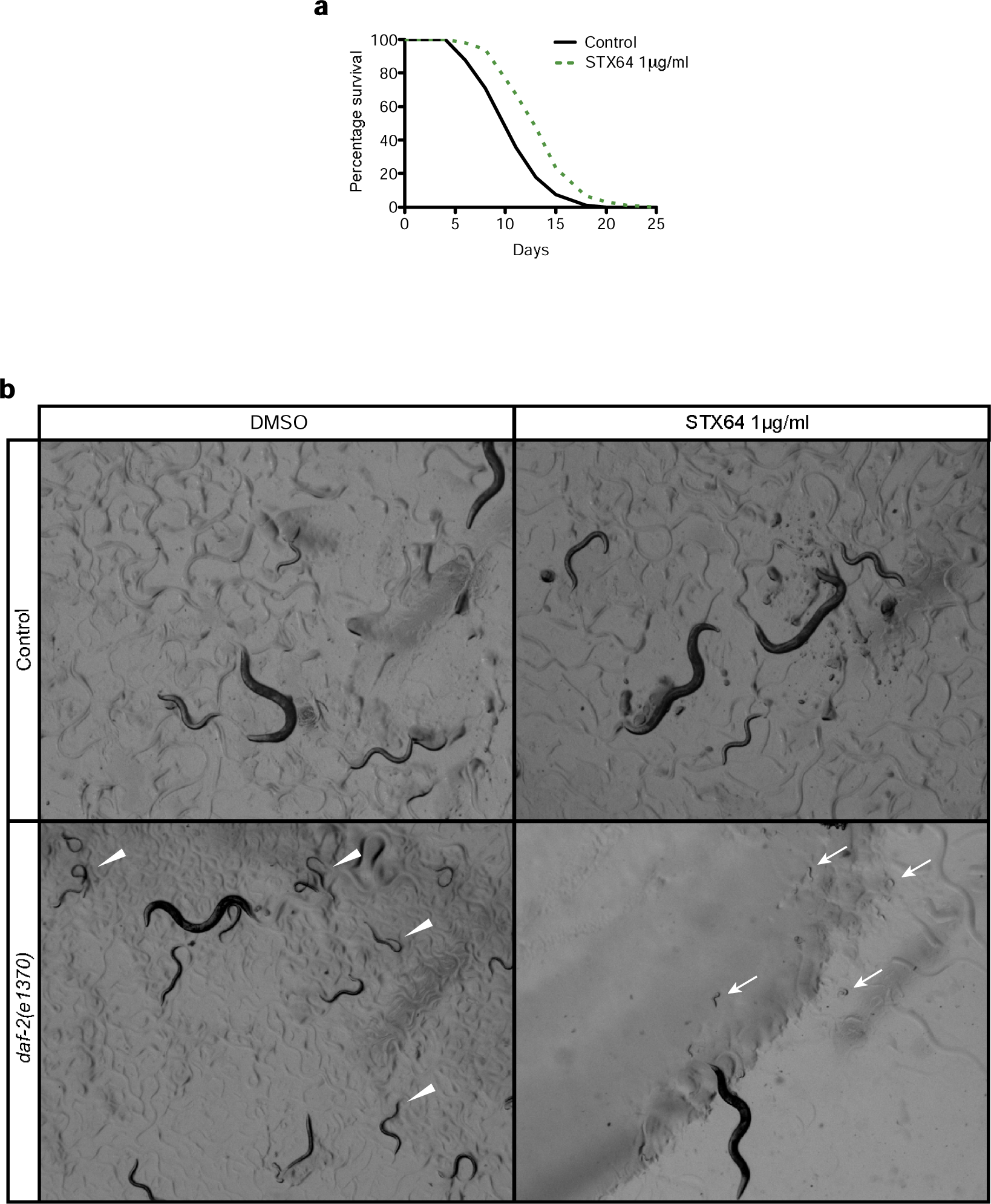
Treatment with STX64 phenocopies longevity and genetic interaction of sul-2 mutants. **a**, STX64 in *non-UV E. coli* increases lifespan of wild type, **b**, Photographs of wild type and *daf-2(e1370)* at 25 °C. DMSO (STX64 vehicle) does not affect development of wild type, neither STX64 treatment, *daf-2(e1370)* arrests mostly in dauer stage at 25°C, but arrests in L1 when treated with STX64, similar interaction is also observed in the *sul-2* mutants. Photographs were taken in a Leica scope. Arrow heads: dauers, arrows: L1s. Statistics of longevity curves are shown in Supplementary Table 1.

**Extended Data Figure 4.**
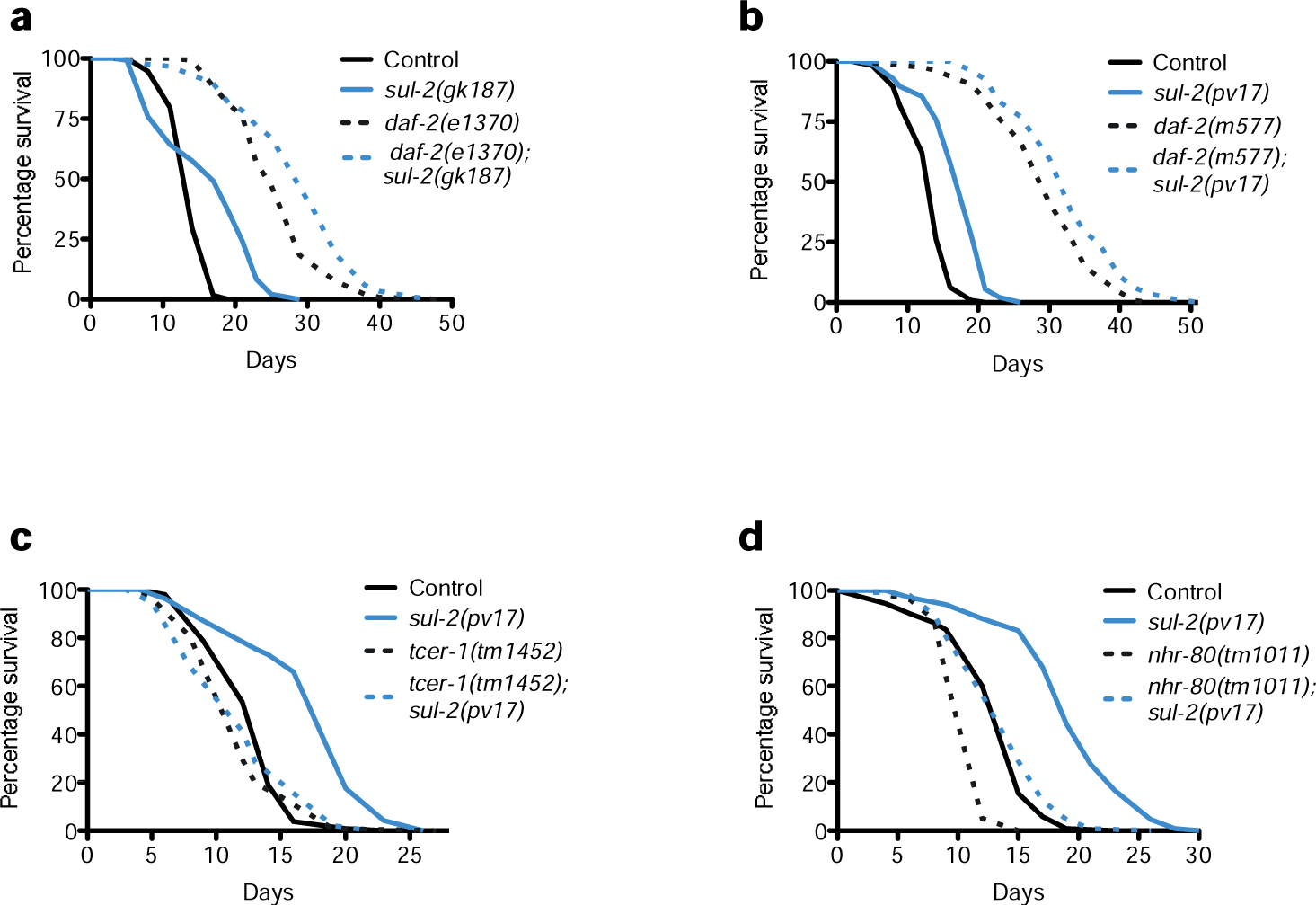
Genetic interactions of *sul-2* mutants. *sul-2* mutants enhace longevity of *daf-2* in non-allele specific way and the longevity is suppressed by mutations in factors involved in the germ line longevity. **a**, *sul-2* deletion mutant enhances longevity of *daf-2(e1370)*. **b**, *sul-2* point mutation enhances longevity of *daf-2(m577)*. **c**, *sul-2(pv17)* longevity is suppresed by *tcer-1(tm1452)*. **d**, *sul-2(pv17)* longevity is suppreses by *nhr-80(tm1011)*. Statistics are shown in Supplementary Table 1.

**Extended Data Figure 5.**
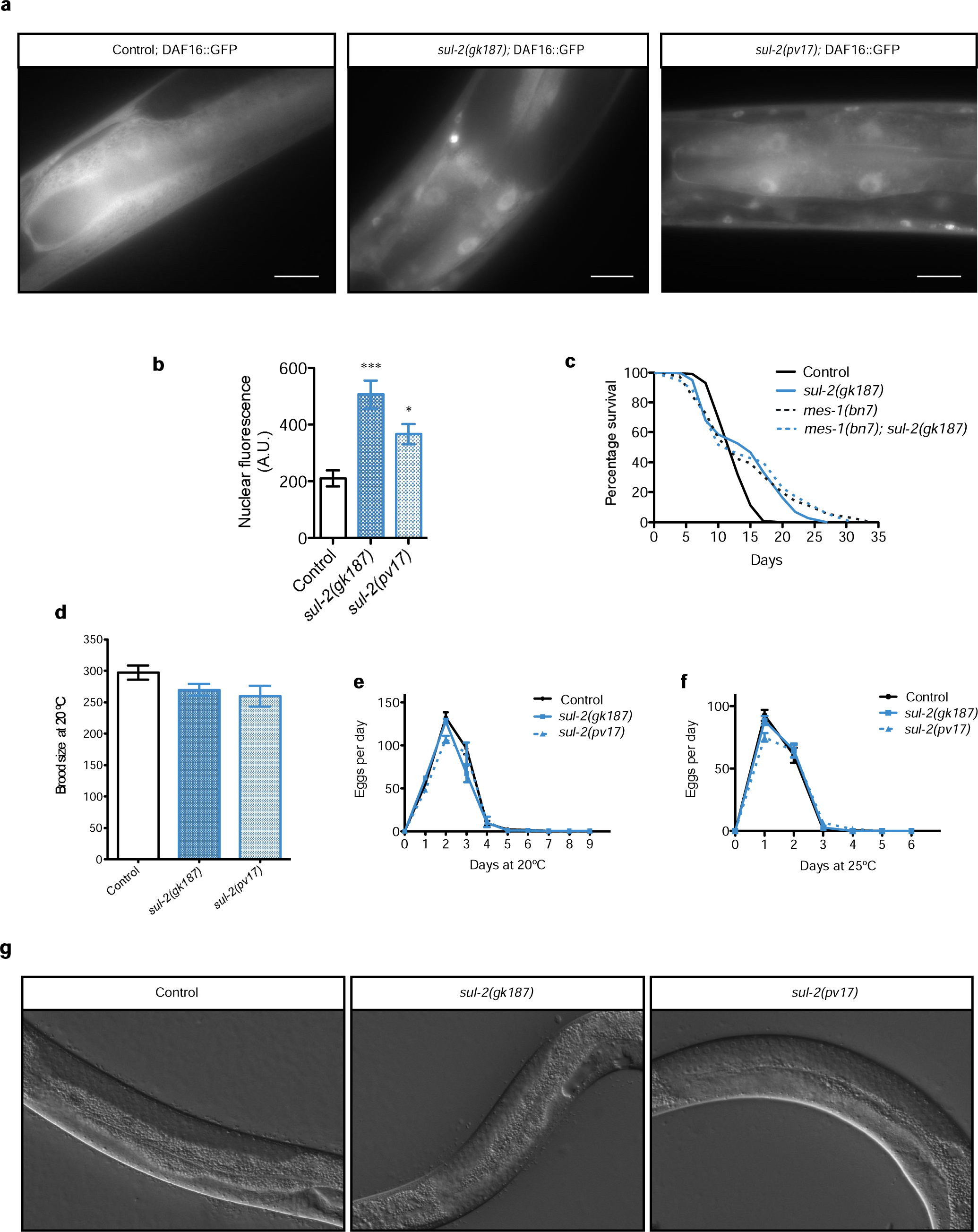
*sul-2* mutants affect DAF-16 location in intestinal cells, but they do not affect reproduction or gonad morphology. **a**, Micrographs show representative images of *Pdaf-16::gfp::daf-16* location in wild type (left panel) and *sul-2* mutants (central and right panels). Both *sul-2* mutants increase the nuclear location of DAF-16 in intestinal cells, like in germ-line less animals. Scale 20 μm. **b**, Quantifications of nuclear fluorescence in the anterior intestinal cells. Data from two independent assays, *n≥* 34 nuclei *per* condition. One-way ANOVA test. **c**, *sul-2* deletion has not significant increase of longevity in germline-less mutant *mes-1(bn7)* mutant background. **d**, *sul-2* mutants have similar brood size to control at 20 °C. One-way ANOVA test; ns. **e**, The reproductive period of *sul-2* mutants are not affected at 20 °C. **f**, Or at 25 °C. **g**, *sul-2* mutants show normal gonad morphology. Micrographs of one representative gonadal arm in late L4 for each strain are shown. Statistics of longevity curves are shown in Supplementary Table 1.

**Extended Data Figure 6.**
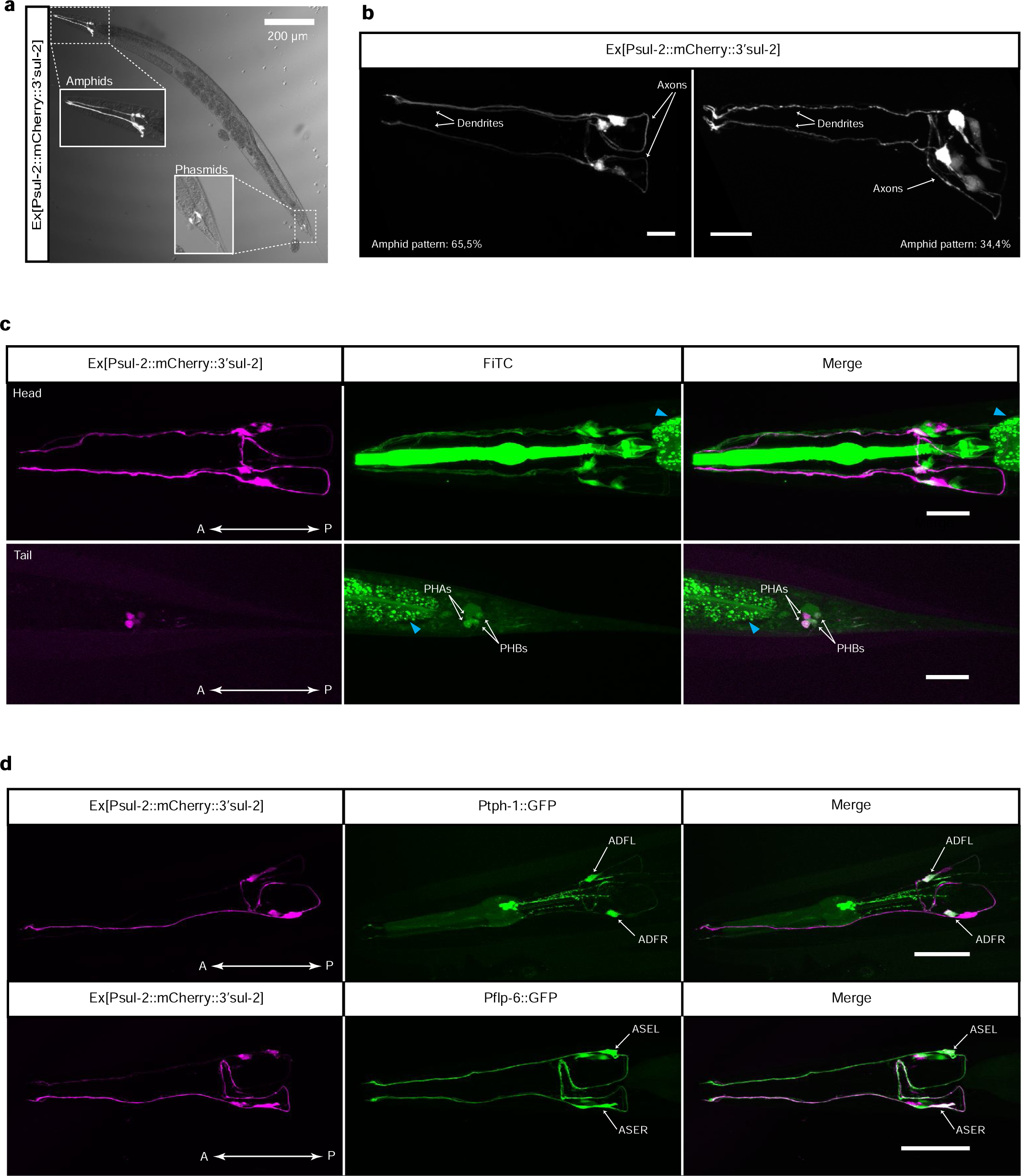
*sul-2* is expressed in amphid and phasmid sensory neurons. Extrachromosomal transgenic worms express mCherry under *sul-2* promoter and its 3’UTR only in few sensory neurons. **a**, Transcriptional reporter for *sul-2* is expressed in sensory neurons. Imaged in fluorescence and merged with the bright field. **b**, Representative images of *sul-2* extrachromosomal expression in amphids, most transgenic animals express *sul-2* in two pairs of amphid neurons, left panel, and a portion show expression in other neurons besides of those, rigth panel. n=64. **c**, Collocation of *sul-2* neurons with FiTC stains. Upper panel shows colocalization in the head with the most anterior pair amphids, possibly ASK, ADF or ADL neurons, but not with the most posterior pair, ASG or ASE. In the tail, bottom panel, the four neurons where *sul-2* is expressed colocalize withFiTC staining in PHAs and PHBs phasmids neurons. **d**, Identification of the two main pair of amphid where *sul-2* is exppresed by collocalization with the *tph-1* neuron-specific promoter for ADF, upper panel, and the posterior neurons expressed by *flp-6* promotor, ASE amphids, bottom panel. Scale bar 20 μm. In intestine, signals are unspecific from autofluorescence at the conditions imaged (cyan arrow heads).

**Extended Data Figure 7.**
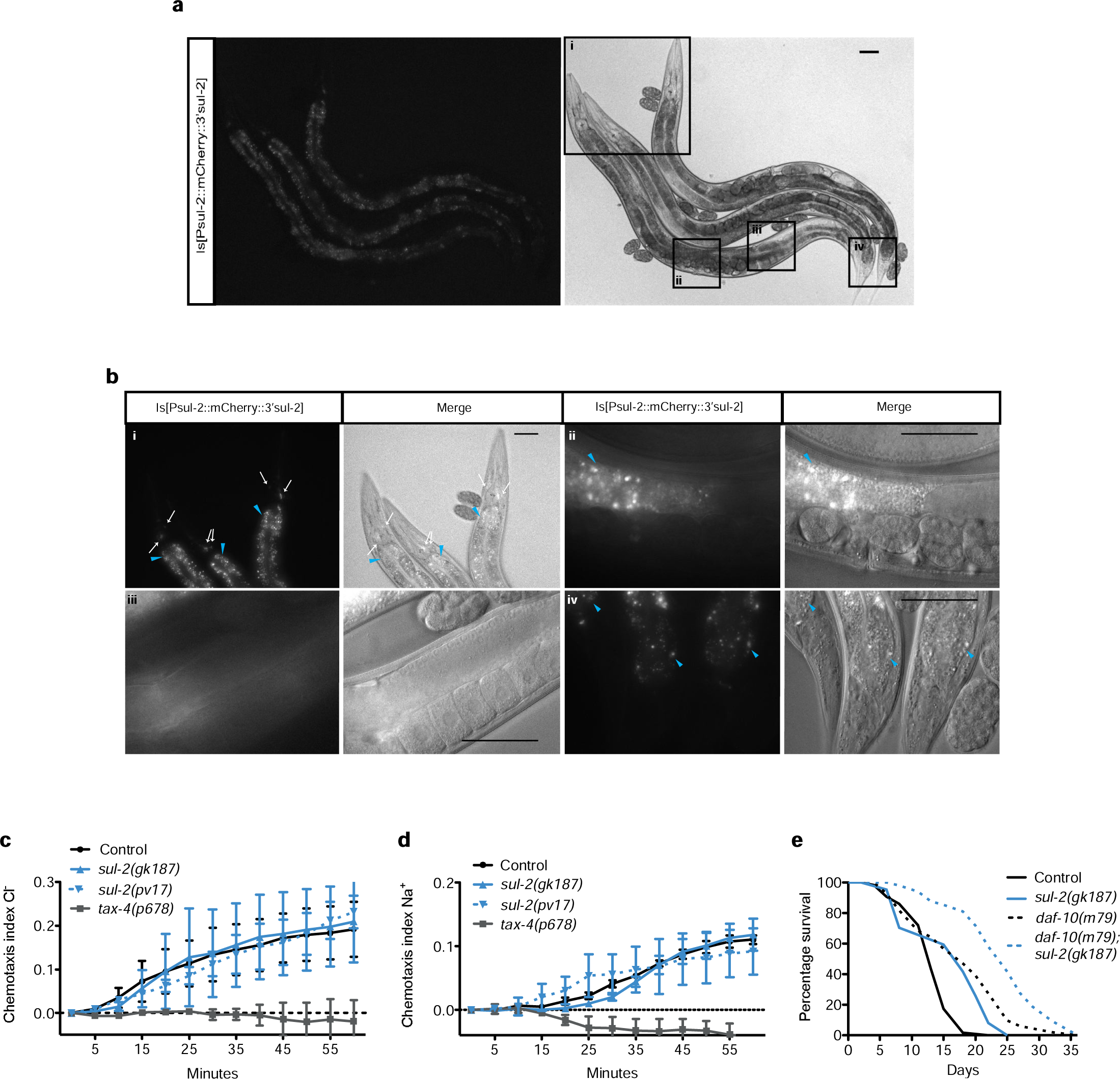
*sul-2* is not expressed in gonadal tissues and is not affected in neuronal functions. **a**, Integrated mCHERRY reporter of the *sul-2* transcriptional unit is only expressed in sensory neurons, there is not expression in other tissues. Image in fluorescence and merged with the bright field image. **b**, Inset of integrated worms, i) In heads, *sul-2* is expressed in few amphid neurons (white arrows). In intestine, signals are unspecific from autofluorescence at the conditions imaged (cyan arrowheads). ii) There is not specific signals in vulva, embryos or proliferative germline zone. iii) There is not specific signals in gonad or mature oocytes. iv) In the tail, there is not significant signals in phasmids. Scale bar **50** μm. **c**, Apart from been expressed in the Cl^−^ and Na+ sensing neuron (ASE), *sul-2* mutants respond to C^−^ similarly to wild type. Data from three independent replicates, **d**, *sul-2* mutants respond to Na+ similarly to wild type. *tax-4(p678)* is a negative control. Data from three independent replicates. In all graphs Mean±SEM are displayed. **e**, *sul-2* deletion enhances longevity of the long-lived *daf-10(m79)* mutant, which is affected in sensory neurons. Statistics of longevity curves are shown in Supplementary Table 1.

**Extended Data Figure 8.**
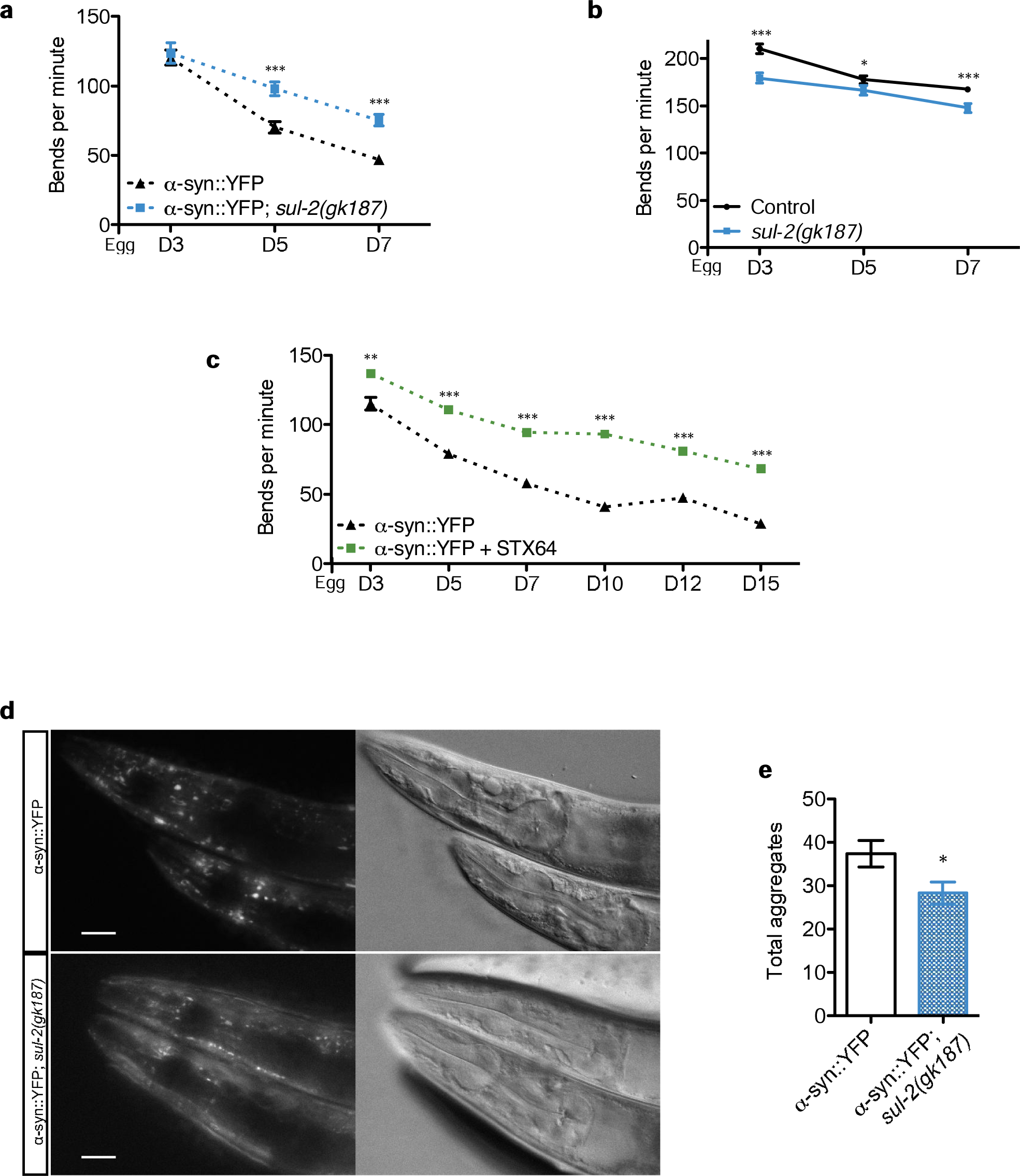
Muscular Parkinson phenotypes are ameliorated during aging when steroid sulfatase function is reduced, and a-synuclein aggregation decreases. The increment of movement is present in all adulthood and is not due to an increase in *sul-2 per se.* **a**, *sul-2* has a beneficial effect during adulthood in muscular Parkinson model. Data display from two independent biological replicates, n≥31 per day and condition, **b**, *sul-2* has less body bends to *wild type* contro, n≥15 per day and condition, **c**, The protective effect of STX64 in muscular Parkinson model is present throughout aging. Data display from two independent biological replicates, n≥12 per day and condition. **d, e**, *sul-2* reduces significantly the number of α-synuclein aggregates in muscle at 7-day old. Photographs exemple of animals and quantifications, respectively. n=18. Scale bar 25 μm.

**Extended Data Figure 9.**
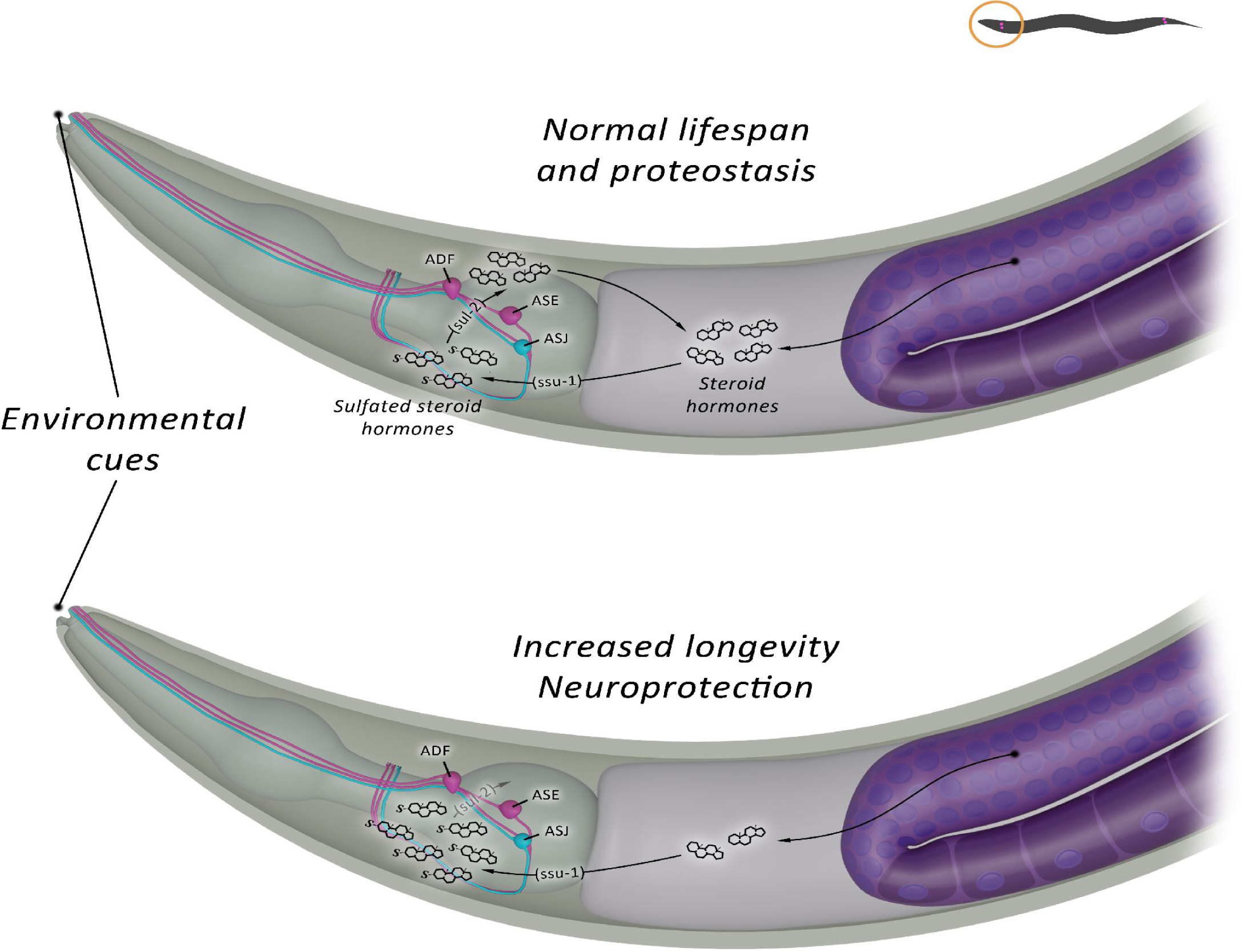
Model of regulation of longevity by SUL-2. Somatic gonads generate a signal to increase longevity through the activation of the nuclear receptor DAF-12. This signal is inhibited by the germline. We propose that one or a pool of unidentified steroid hormones mediate this inhibiting signaling which remarkably is independent of the reproductive function. This steroid hormone can be inactivated by sulfation and activated back by the activity of the steroid sulfatase SUL-2, although we can not rule out the possibility that sulfated steroid hormones can have a prolongevity activity. The fact that *sul-2* is expressed in sensory neurons suggests that the sulfated state of these steroid hormones can be regulated by environmental signals. The reduction of *sul-2* activity either by mutation or by STX64 treatment unbalances the level of sulfated hormones, probably reducing level of non-sulfated hormones and increasing sulfated hormones, this results in an increase of longevity and also an increase level of proteostasis, not only in *C. elegans* but also in mammals.

